# Aged brain multi-omic integration captures immunometabolic and sex variation

**DOI:** 10.1101/2025.10.10.681695

**Authors:** Justin P. Whalley, Holly C. Hunsberger, David A. Bennett, Melissa Lamar, Daniel A. Peterson, Grace E. Stutzmann

## Abstract

Resolving the molecular basis for heterogeneous ageing in the human brain requires integrating its diverse molecular layers. Here we applied a tensor-decomposition framework to jointly analyse single-nucleus RNA-seq, DNA methylation, H3K9ac histone acetylation and proteomic data from 276 post-mortem human cortices. The analysis uncovered two dominant, multi-omic themes. First, we identified an immunometabolic axis characterised by microglial activation, suppression of PI3K-Akt-mTOR signalling and epigenetic signatures that suggest repurposing of developmental transcription factors such as BARHL1. Second, we decomposed the effect of sex into a multi-omic program linked to cellular plasticity: although driven by canonical sex-chromosome genes, this axis was epigenetically enriched for pathways regulating stem-cell pluripotency and exhibited a female-biased shift toward precursor-like states in the oligodendrocyte lineage. Together, these findings deliver a high-resolution atlas of glial ageing and demonstrate that broad risk factors manifest as complex, cell-specific vulnerabilities; an insight critical for the development of targeted therapeutic strategies.

## INTRODUCTION

Aging of the human brain, and the development of neurodegenerative diseases such as Alzheimer’s disease (AD), is accompanied by intertwined alterations in gene expression, epigenetic regulation, and protein homeostasis that unfold across diverse cell types (Iturria-Medina et al., 2022). Recent large-scale initiatives have generated richly annotated, multi-omic resources from post-mortem human cortex, most notably the Religious Orders Study and Memory and Aging Project (ROSMAP) (Bennett et al., 2018). These datasets create an unprecedented opportunity to integrate transcriptomic, epigenomic and proteomic layers and obtain a systems-level view of cognitive decline and dementia progression.

However, these omic layers are often studied in isolation, leaving the complex interplay between them poorly understood. A key challenge is to move beyond correlational studies and develop a more causal understanding of how changes in the epigenome, for example, propagate to alter transcription and protein function within specific cell types (Gabitto et al., 2024). To meet this challenge, computational frameworks capable of jointly modeling heterogeneous data are required. Tensor decomposition methods are particularly well-suited for this task, as they can deconstruct multi-modal datasets into a set of coherent, underlying biological components or programs. We and others have successfully used this approach to uncover systems-level patterns in other complex human diseases, demonstrating its power to find novel biological insights amidst high-dimensional data (COvid-19 Multi-omics Blood ATlas (COMBAT) Consortium, 2022; Mi et al., 2024).

This integrative approach is critically needed to dissect the roles of glia in brain aging. Microglia and astrocytes, once seen as passive support cells, are now recognized as central drivers of neuroinflammation and metabolic reprogramming in AD (De Strooper & Karran, 2016). Epidemiologically, women are at higher risk of developing AD, yet the cellular and molecular basis for this vulnerability remains poorly defined (Buckley et al., 2023). Understanding how sex shapes glial states at the transcriptional and epigenetic levels is a major gap in the field (Casaletto et al., 2022). Underlying these glial states are fundamental homeostatic processes, including the maintenance of myelin by oligodendrocytes (Behrendt et al., 2013; Perlman et al., 2020; Adams et al., 2021) and the precise regulation of intracellular calcium signaling (Stutzmann, 2007; Mustaly-Kalimi et al., 2022, 2025), both of which are known to be dysregulated in AD.

Here, using integrative analysis of transcriptomic, epigenomic and proteomic data, we reveal two dominant features of the aged brain. First, we characterize a major immunometabolic axis driven by microglial activation, the suppression of PI3K-Akt-mTOR signaling, and the apparent repurposing of developmental epigenetic programs. Second, we resolve the broad effect of sex into a mosaic of distinct, cell-type-specific molecular programs within glia, including male-biased mitochondrial programs in microglia and a shift toward precursor-like states in the female oligodendrocyte lineage. This work provides a high-resolution map of how risk factors like age and sex manifest as complex, cell-specific vulnerabilities.

## RESULTS

We aimed to explain variation in the aging brain through integration of multi-omic data from the Religious Orders Study and Memory and Aging Project (ROSMAP) (Bennett et al., 2018). Our primary dataset comprised single-nucleus RNA sequencing (snRNA-seq) from 276 prefrontal cortex samples, ensuring a minimum of 100 cells per subtype across astrocytes, oligodendrocytes, inhibitory neurons, oligodendrocyte precursor cells (OPCs), excitatory neurons, and microglia (Mathys et al., 2023). This transcriptomic data was complemented by three additional omic layers: DNA methylation at 51,798 CpG sites from (216 dorsolateral prefrontal cortex samples) (De Jager et al., 2014), chromatin immunoprecipitation sequencing (ChIP-seq) of H3K9ac histone marks at 26,384 peaks (193 prefrontal cortex samples) (Klein et al., 2019), and mass spectrometry proteomics quantifying 5,230 proteins (105 dorsolateral prefrontal cortex samples) (Ping et al., 2018). The integration was anchored on the 276 samples with snRNA-seq data, as this cohort had the largest sample size and provided the cellular resolution necessary for our analysis. This approach maximized sample overlap across all four omic types.

To unify these heterogeneous molecular layers, we applied Sparse Decomposition of Arrays (SDA), a Bayesian tensor factorization method that we adapted for this research (Hore et al., 2016). We chose SDA for its ability to reduce dimensionality across all four modalities and handle missing data, which is critical for complex biological systems. Our decomposition of this integrated dataset resulted in 546 latent components, each capturing a coherent axis of biological variance across omic types (Figure 1). This approach allowed us to resolve both cell-specific programs and cross-modal regulatory circuits.

**Figure 1.**
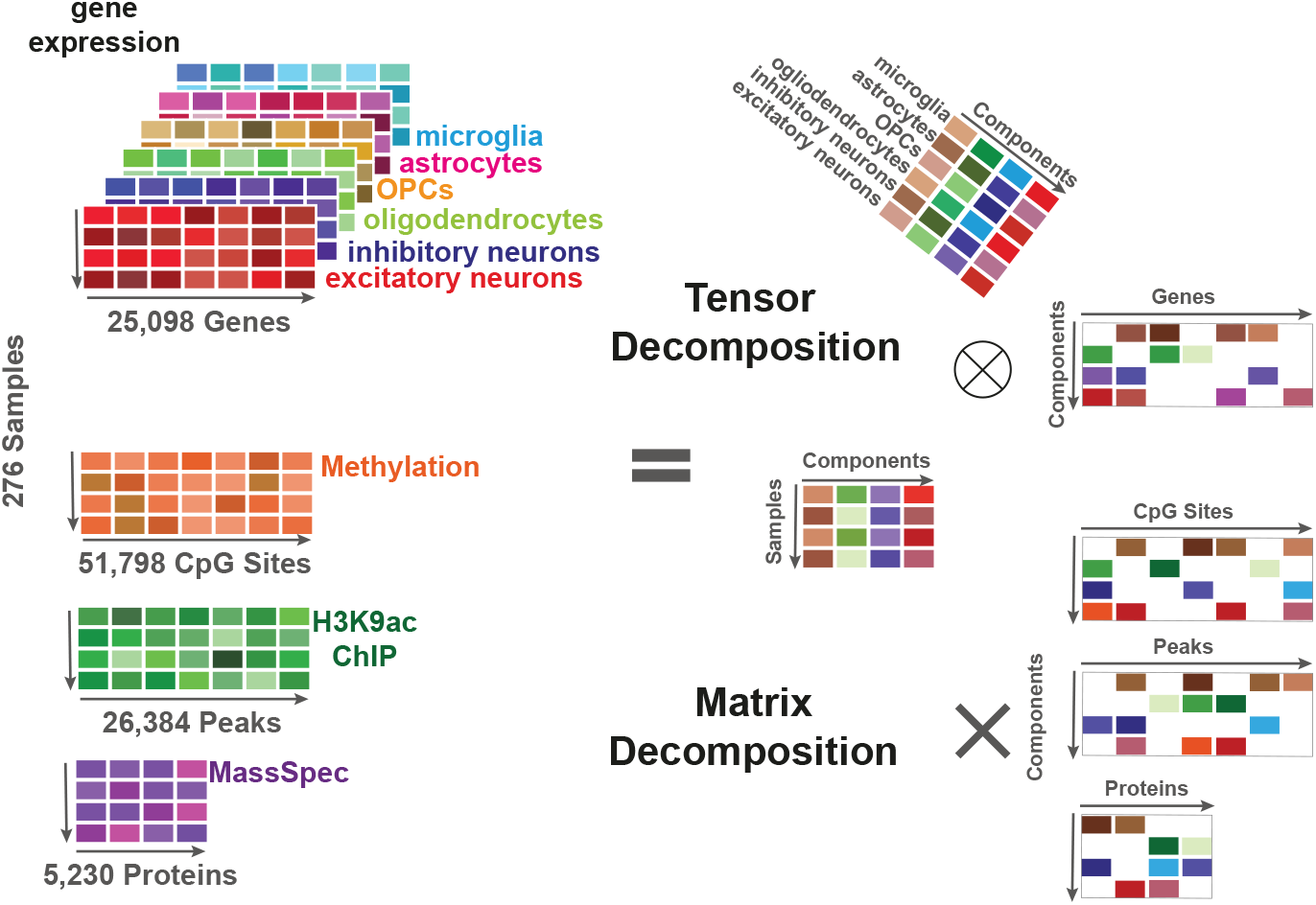
Schematic of work: tensor and matrix decomposition applied to the ROSMAP datasets. The left panel shows the input data: a three-dimensional gene expression tensor (samples × genes × cell types) and three two-dimensional matrices for DNA methylation (CpG sites), histone acetylation (ChIP-seq peaks), and proteomics (proteins), all aligned by sample. The right panel depicts the output of integrative decomposition using Sparse Decomposition of Arrays (SDA), which yields a shared set of latent components. Each component is represented by a vector of sample loadings, combined via tensor multiplication with cell type and gene loadings for the RNA-seq data, and via matrix multiplication with sparse feature loadings for the methylation, ChIP-seq, and proteomic data.

As an initial validation of the method’s ability to uncover novel biological signals within our data, SDA revealed a previously unreported eQTL regulating a cluster of olfactory receptor genes, with the signal being driven predominantly by neuronal and oligodendroglial cells (Component 21 as shown in Figure S1). This finding not only validates our analytical approach but also highlights the complex regulatory landscape of ectopic gene expression in the brain, as olfactory receptors have been implicated in diverse cellular processes beyond olfaction (Maßberg & Hatt, 2018).

From the 546 components, we first generated a comprehensive annotation, summarized in Supplementary Table 1, that details each component’s primary cell-type drivers, top multi-omic features, pathway enrichments, and significant associations with clinical phenotypes (such as cognitive decline and biological sex). Guided by this overview, we then prioritized a subset of components for detailed characterization in the main text based on their strong multi-omic coherence and robust cross-cohort reproducibility in the SEA-AD snRNA-seq dataset from 82 dorsolateral prefrontal cortex samples (Gabitto et al., 2024). The following sections are organized around two key themes that emerged from this filtered set: the dissection of complex immunometabolic axes within glial cells, and the resolution of sex differences into multiple, distinct, and cell-type-specific signatures. All component loading scores and the code used for this analysis are publicly available on Synapse and GitHub, respectively.

### Component 351: A microglial immunometabolic axis in the aged brain

Integrative tensor decomposition of our multi-omic dataset revealed an immunometabolic program captured by Component 351. Figure 2A displays the loading scores for each major cell type in the snRNA-seq data. Notably, microglial transcripts associated with immune-inflammatory responses, such as *IGKC, IGHG1, CD68*, and *HLA-DRB1*, show strong contributions to this component (Figure 2B), suggesting robust microglial activation in the aged brain (Figure S2A).

**Figure 2.**
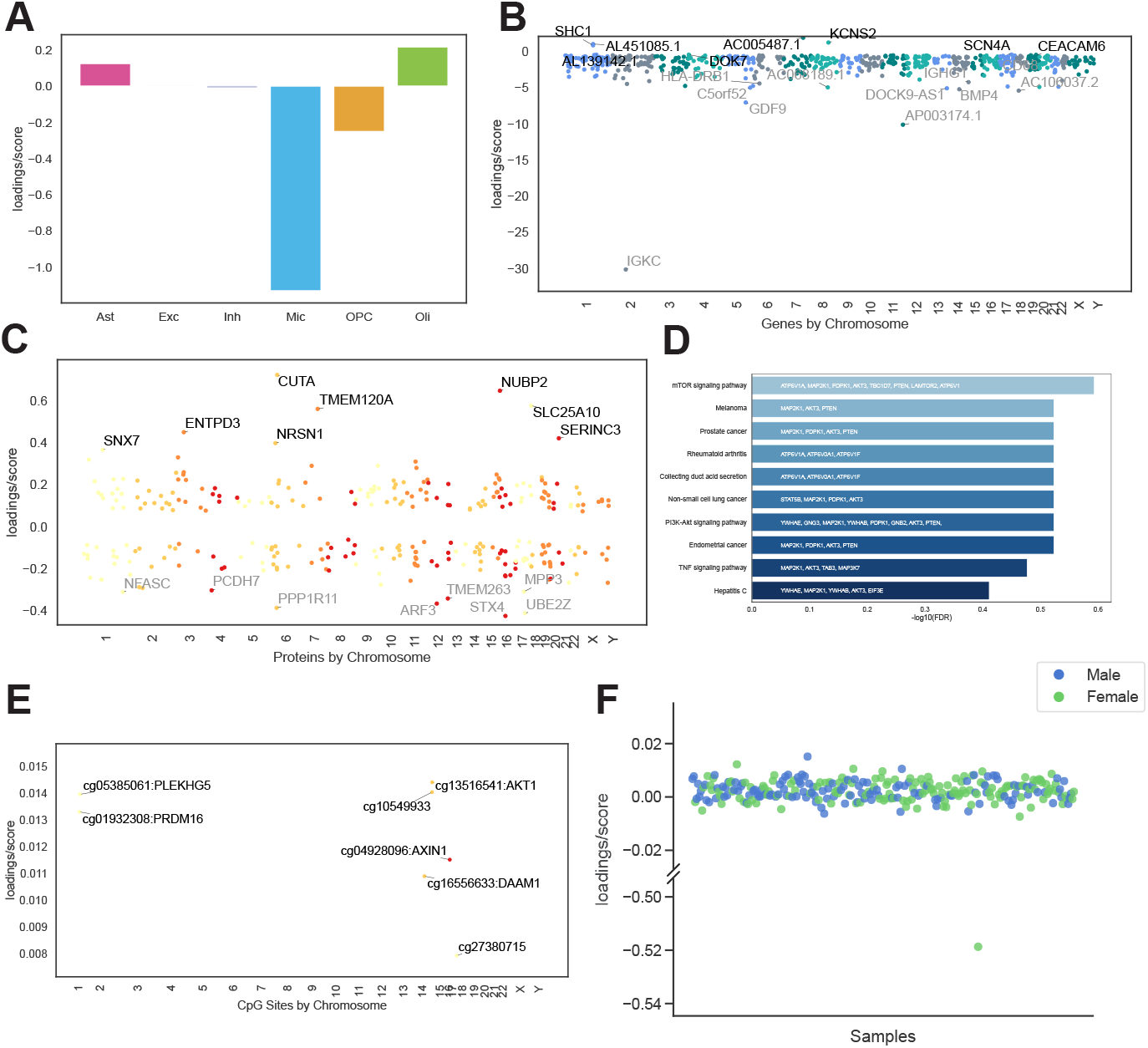
A microglial immunometabolic program in the aged brain (Component 351). **(A)** Cell-type loadings highlight strong microglial contributions to this component. **(B)** Top gene loadings from snRNA-seq show elevated expression of immunoglobulin and antigen-presentation transcripts (e.g., *IGKC, IGHG1, CD68, HLA-DRB1*), consistent with immune activation. **(C)** Positively loaded proteins include antioxidant and stress response factors (e.g., CUTA, SERINC3, SEP15), while negatively loaded proteins reflect downregulation of PI3K-Akt-mTOR signaling (e.g., AKT3, PTEN). **(D)** KEGG pathway enrichment for negatively loaded proteins shows suppression of metabolic and growth pathways, including mTOR, PI3K-Akt, and TNF signaling. **(E)** CpG sites near immune and metabolic genes (e.g., *AKT1, DAAM1, PLEKHG5*) show consistent loading directions, supporting epigenetic regulation of this axis. **(F)** Sample-level scores across individuals, with one outlier showing markedly high loading score, potentially indicating a localized or atypical inflammatory process.

Protein-level analysis (Figure 2C) reinforces the same pro-inflammatory direction, featuring multiple transporters and antioxidant proteins (e.g., *CUTA, SERINC3, SEP15*) that may reflect metabolic reprogramming under immune stress. In contrast, the subset of proteins exhibiting negative loadings (Figure 2D) is significantly enriched for PI3K-Akt-mTOR pathway members (e.g., *AKT3, PTEN, PDPK1, MAP2K1*), implying that microglial activation could coincide with suppressed growth signaling or cell-type loss within these aged samples.

Methylation profiling (Figure 2E) adds a further dimension, highlighting CpG loci near *AKT1* and other metabolic or immune genes that load in the same pro-inflammatory, low-mTOR direction. Additional evidence comes from H3K9ac ChIP-seq peaks (Figure S2B) enriched for a *BARHL1*-like motif. Although *BARHL1* is best known for its role in neurodevelopment and cerebellar formation (Lopes et al., 2006), our data imply a possible re-engagement of this transcriptional network, in older brains under high inflammatory states, raising questions about BARHL1’s function or its related homeobox factors in chronic neuroinflammation or neural remodeling. The re-engagement of such a developmental transcription factor in the aged brain could signify a maladaptive or incomplete attempt at neural remodeling or a cellular identity shift in response to persistent immune stress.

Parallel analysis in the independent SEA-AD dataset revealed a correlated transcriptional signature (Component 21) that marks a principal immune axis in aged cortex (Figure S2C-E). This correspondence validates the microglial-immune program captured by ROSMAP Component 351. In both cohorts, a few outlier samples display exceptionally high *IGKC* expression (Figure 2F; Figure S2E), consistent with sporadic B-cell-driven inflammation. Importantly, although SEA-AD Component 21 appears as a single transcriptional component, our multi-omic decomposition in ROSMAP resolves it into two discrete biological programs: Component 351, an immunometabolic axis, and Component 2, an immune-regulatory axis discussed next.

### Component 2: A glial immune-regulatory axis balancing inflammation and homeostasis

Component 2 is characterized transcriptionally by a strong microglial signature whose leading feature is the immunoglobulin gene *IGKC* (Figure 3A-B). This signature is also correlated with SEA-AD Component 21 (Figure S3B). Unlike Component 351, however, Component 2 is defined by unique multi-omic features and cell-type drivers: its hallmark is a pronounced inverse relationship between microglial/B-cell activation and astrocytic programs (Figure 3A). Thus, Component 2 reflects a reciprocal glial state in which immune activity rises while astrocytic pathways are selectively suppressed, and vice versa, rather than uniform glial activation.

**Figure 3.**
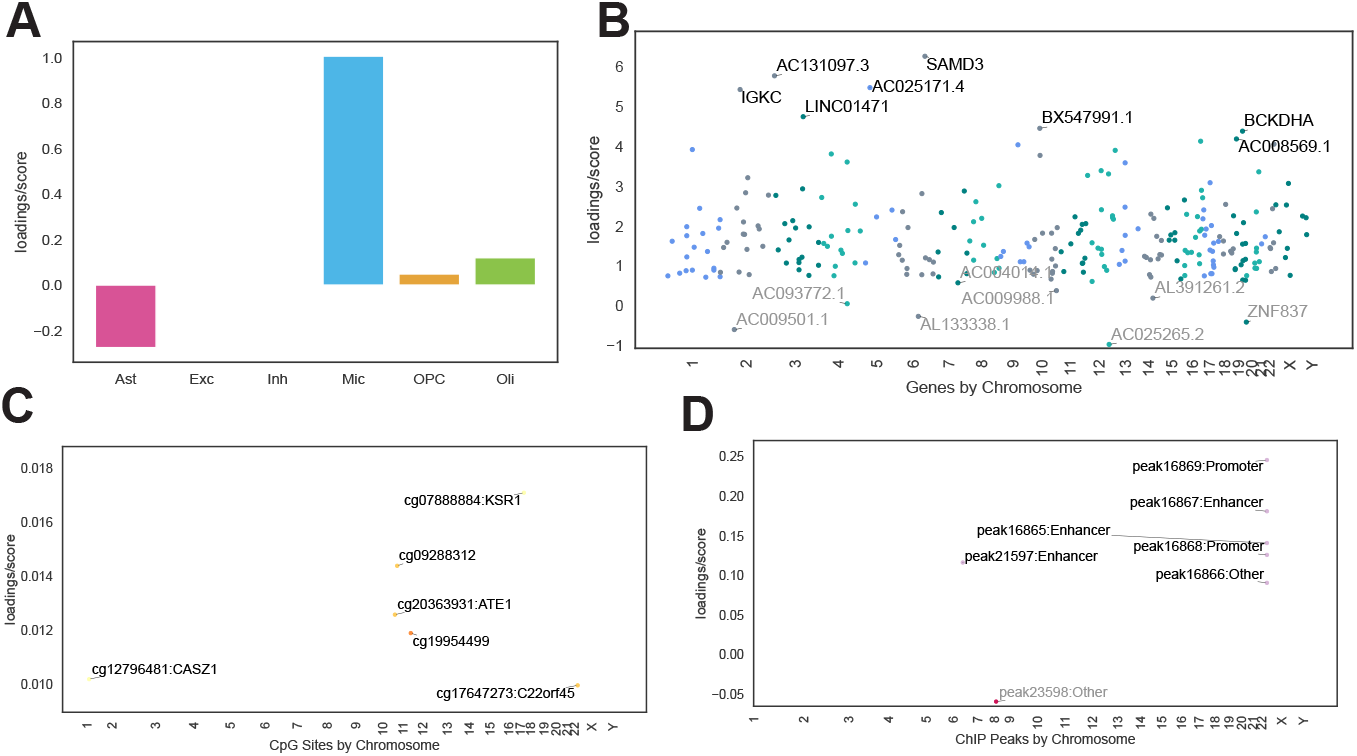
Multi-omic features and regulatory annotations for Component 2 in the aging brain. **(A)** Cell-type loadings from snRNA-seq data, highlighting positive microglial and negative astrocytic contributions to Component 2. **(B)** Top gene loadings from snRNA-seq, highlighting non-coding transcripts among the top loadings. **(C)** Methylation sites with high PIP values for Component 2, annotated to genes including *KSR1, CASZ1*, and *ADORA2A-AS1*. **(D)** H3K9ac ChIP-seq peak loadings, showing a cluster of enhancer regions on chromosome 22 overlapping the *CACNA1I* and *ENTHD1* locus. These epigenomic signals support the role of purinergic signaling in this glial axis.

DNA methylation patterns add a further layer to this glial axis. Several CpG sites (e.g., cg07888884 near *KSR1*, cg12796481 near *CASZ1*, and cg17647273 in *ADORA2A-AS1*, labelled as C22orf45) are differentially methylated in association with Component 2 loadings (Figure 3C). These loci map to signaling and metabolic regulators implicated in glial reactivity and inflammation resolution. Chromatin accessibility data (ChIP-seq) reveal a cluster of active enhancer peaks on chromosome 22 (Figure 3D), overlapping *CACNA1I*, which encodes a T-type voltage-gated calcium channel, and *ENTHD1*. Calcium signaling is a fundamental mechanism of communication for both neurons and glial cells, suggesting that immunoglobulin-associated signal is functionally coupled to alterations in the brain’s core calcium signaling machinery.

Together, these features suggest that Component 2 captures a dynamic glial state characterized by immune engagement from microglia and either compensatory suppression or reactivity in astrocytes. This is further supported by the prevalence of uncharacterized or non-coding transcripts among top RNA-seq loadings, points to variation in how aging brains manage immune stress (Figure 3B). In parallel to Component 351, which reflects a coordinated upregulation of immune and metabolic genes, Component 2 appears to mark an intrinsic regulatory balance between glial cell types, pointing to variation in how aging brains manage immune stress.

### Component 74: Resolving sex differences in neurons and glial lineages

Our decomposition resolved multiple axes of sex-biased molecular variation, the most significant of which was Component 74 (two-sided Mann-Whitney; *U* = 18962, *p* = 1.5 × 10^−46^, FDR-adjusted *p* = 7.9 × 10^−44^) (Figure 4A). This component captured the canonical sex signature, with gene loadings dominated by X- and Y-chromosome transcripts (e.g., *XIST, UTY*) (Figure 4D). Transcriptionally, this program was strongest in excitatory and inhibitory neurons, but interestingly, showed a strong opposing effect in oligodendrocytes (Figure 4B). This core signature was robustly validated, showing a inverse correlation (*r* = −0.982) with the primary sex component in the independent SEA-AD dataset (Figure 4C).

**Figure 4.**
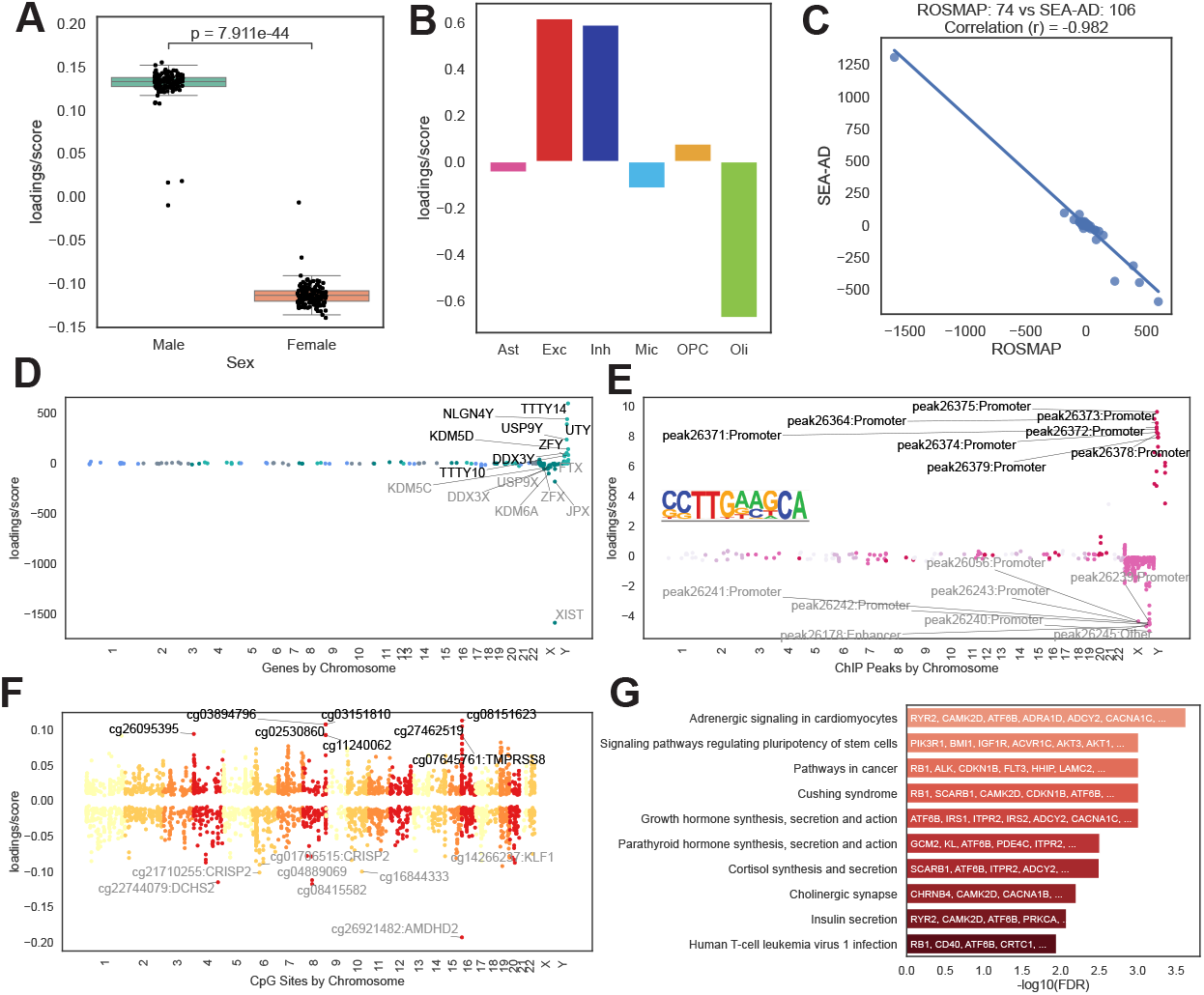
Sex-difference in neuronal cells as shown in component 74. **(A)** Box plots (median ± IQR; whiskers extend to 1.5 × IQR) of sample scores for Component 74, showing significant separation between male and female individuals (FDR-adjusted *p* = 7.9 × 10^−44^). **(B)** Cell-type loadings, showing strong positive contributions from excitatory (Exc) and inhibitory (Inh) neurons, contrasted by a strong negative contribution from oligodendrocytes (Oli). **(C)** Scatter plot showing near-perfect anti-correlation (*r* = −0.982) between ROSMAP Component 74 and the corresponding sex component in the SEA-AD dataset. **(D)** Gene expression loadings are dominated by canonical X- and Y-chromosome genes. **(E)** H3K9ac ChIP-seq peak loadings are concentrated on the sex chromosomes and are enriched for a binding motif of the stem-cell-related transcription factor *PRDM4*. **(F)** DNA methylation loadings show widespread, sex-biased differences across all autosomes. **(G)** KEGG pathway enrichment for genes near sex-differentiated CpG sites.

Beyond these canonical features, Component 74 revealed a surprisingly deep, multi-omic signature of sex that implicates the regulation of cellular plasticity. We observed widespread, sex-differentiated DNA methylation patterns across the autosomes, not just the sex chromosomes (Figure 4F). Pathway analysis of the genes associated with these CpG sites revealed “Signaling pathways regulating pluripotency of stem cells” as one of the top enriched KEGG pathways (Figure 4G). This finding was reinforced by the chromatin accessibility data. The component’s ChIP-seq signature was epigenetically marked by promoter peaks on the X and Y chromosomes, which were enriched for binding motifs of PRDM4: a transcription factor known to play a key role in stem cell self-renewal (Figure 4E). Together, these results demonstrate that the primary axis of sex difference in the aging brain is not merely a reflection of sex chromosome gene dosage, but a complex, multi-layer program tied to the fundamental regulation of cellular plasticity and stem-like states.

This theme of sex-differentiated cellular plasticity is further elaborated by Component 278, which focuses specifically on the oligodendrocyte lineage. Sample scores for this component were significantly higher in females (two-sided Mann-Whitney; *U* = 6808, *p* = 5.3 × 10^−5^, FDR-adjusted *p* = 0.0097) (Supplementary Figure 4A). The cell-type loadings reveal a dynamic axis of opposition between oligodendrocyte precursor cells (OPCs), which contribute positively, and mature oligodendrocytes, which contribute negatively (Supplementary Figure 4B). This suggests that a higher score, as seen more frequently in females, reflects a cellular state that is transcriptionally closer to an OPC/precursor state and less mature. The biological basis of this axis is underscored by the enrichment of genes involved in canonical growth and differentiation pathways, including RAS, PI3K-Akt and MAPK signaling (Supplementary Figure 4E), providing a cell-type-specific example of the broader pluripotency theme identified in Component 74.

### Component 87: Glial variation beyond sex and immunity

Beyond the major immunometabolic and sex-driven programs, our analysis resolved further axes of glial heterogeneity defined by robust multi-omic signatures. Here we highlight two key components: a complex astrocyte-OPC regulatory program (Component 87) and a microglial bioenergetic program (Component 177).

Component 87 captures a complex multi-glial program characterised by an opposing relationship between astrocytes and oligodendrocyte precursor cells (OPCs) (Figure 5A). The transcriptional signature of this component is enriched for immunomodulatory genes, with a top hit being *VTCN1* (also known as *B7-H4*), a key negative regulator of T-cell responses (Figure 5B). This immunomodulatory theme is supported by a coordinated program of epigenetic changes. In the DNA methylation data, the component is associated with opposing methylation patterns at CpG sites within the neuronal calcium channel gene *CACNA1C* and near *POLR1A*, a core component of the RNA polymerase I machinery (Figure 5C). This theme is echoed in the chromatin accessibility data, where the component is linked to decreased accessibility at specific promoter peaks (Figure 5D), including those for the key glial regulatory kinases *AXL* and *FLT4*.

**Figure 5.**
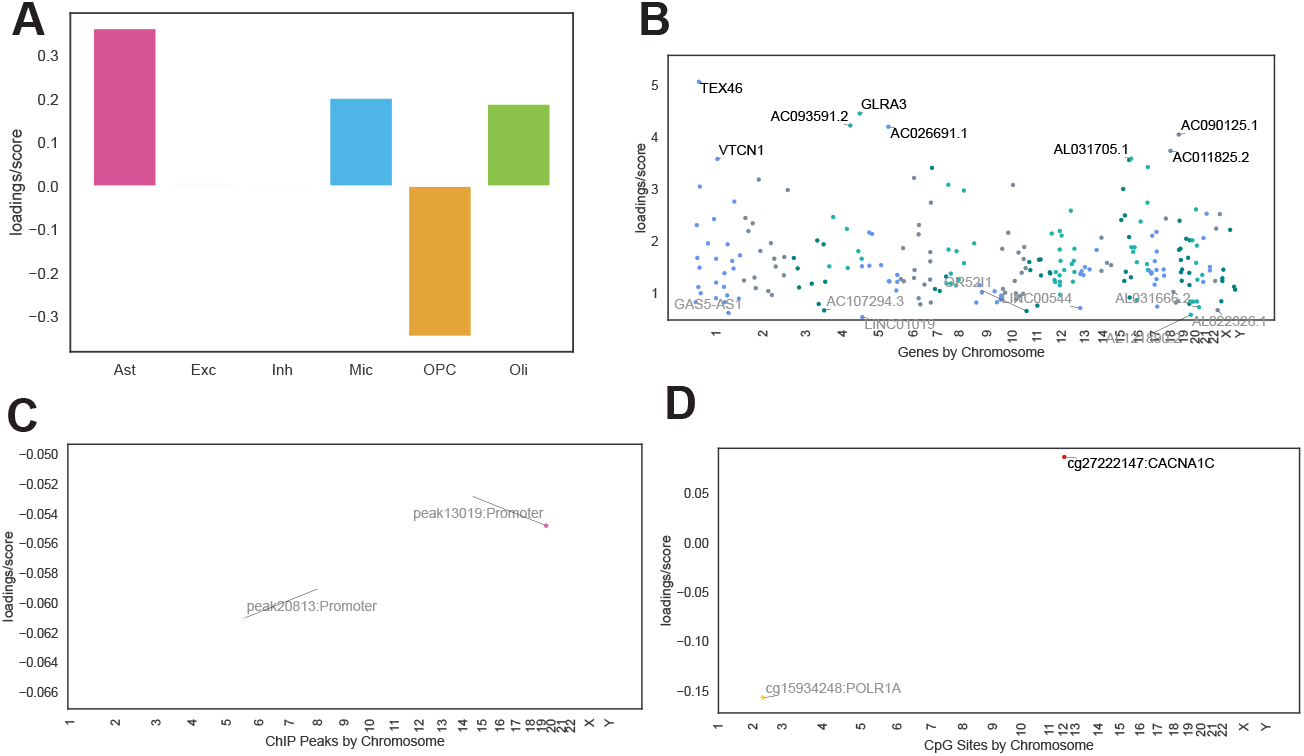
Astrocyte-driven component links immune modulation to coordinated epigenetic reprogramming. **(A)** Cell-type loading scores for Component 87, highlighting a strong positive contribution from astrocytes (Ast) and an opposing negative contribution from oligodendrocyte precursor cells (OPC). **(B)** Gene expression loadings for Component 87, with top hits including the immunomodulatory gene *VTCN1*. **(C)** DNA methylation loadings for Component 87, showing opposing regulation between a CpG site in *CACNA1C* (positive loading) and one near *POLR1A* (negative loading). **(D)** ChIP-seq peak loadings for Component 87, showing decreased accessibility at specific promoter sites.

A second distinct axis of glial variation is captured by Component 177, a program driven almost exclusively by microglia (Supplementary Figure 5A). The transcriptional signature of this component is defined by a coherent set of genes related to mitochondrial bioenergetics (Supplementary Figure 5B), with top hits including *NDUFA13* (a core subunit of Complex I) and *COX6A1* (a subunit of Complex IV). This bioenergetic theme is reinforced at the protein level, where the component’s top hit is Metadherin (MTDH), a protein implicated in cell metabolism and stress (Supplementary Figure 5C). This component’s multi-omic signature is further defined by a coordinated program of DNA methylation changes linked to cellular quality control, including sites near the autophagy initiator ATG9B, the lysosomal enzyme *CTSZ*, and the oppositely regulated cell adhesion gene *ITGA7* (Supplementary Figure 5D). Together, this component appears to capture a microglial state characterized by the joint regulation of mitochondrial energy production and autophagy–lysosomal quality-control pathways.

## DISCUSSION

Our integrative multi-omics analysis reveals previously unappreciated molecular programs that emerge when transcriptomic, epigenomic, and proteomic layers from the aging human cortex are examined in concert. Using Sparse Decomposition of Arrays (SDA), we distilled these layers into two overarching themes: a glial immuno-metabolic axis driven by microglial activation, and a mosaic of pervasive, cell-type-specific sex differences across glial lineages. These findings provide a high-resolution map of how risk factors like age and sex manifest as complex, cell-specific vulnerabilities that are coordinated across multiple biological layers.

A feature of the aged brain was a tightly coordinated immunometabolic program linking pro-inflammatory gene and protein expression with the suppression of PI3K-Akt-mTOR signaling. This coupling supports the concept of inflammaging, where persistent, low-grade immune activity and metabolic dysregulation create a self-reinforcing loop (Franceschi et al., 2018). Interestingly, our finding of suppressed mTOR signaling in the human cortex presents a nuanced contrast to studies in aging female mice, which report heightened mTOR signaling as a driver of microglial glycolysis (Kang et al., 2024). This apparent discrepancy may reflect fundamental differences between species, brain regions, or, perhaps most likely, the distinction between adaptive metabolic shifts in aging versus the chronic, potentially exhausted or maladaptive state of glia in the context of human neurodegenerative pathology.

An unexpected feature of this immunometabolic axis was the enrichment of BARHL1-like motifs in H3K9ac-marked chromatin. As BARHL1 is typically restricted to neurodevelopment, its re-emergence suggests that under prolonged stress, aging glia may re-engage latent developmental programs. This convergence of immune and developmental pathways aligns with a growing understanding of cellular plasticity in disease, where cells can adopt dysfunctional or hybrid identity states (De Strooper & Karran, 2016). This raises the possibility that targeting the epigenetic machinery that governs these latent programs could be a novel strategy to reset maladaptive glial states in late life.

Beyond immunometabolism, our analysis provides a high-resolution view of how sex shapes the brain’s molecular landscape. The canonical sex component (Component 74), driven by X- and Y-chromosome genes, revealed a deeper multi-omic signature linked to the regulation of cellular plasticity. Genes near sex-differentiated CpG sites were enriched for “Signaling pathways regulating pluripotency of stem cells,” and the component’s chromatin signature was marked by motifs for PRDM4. This transcription factor is a key regulator of neural stem cell proliferation and differentiation and also interacts with the PI3K/AKT pathway, providing a fascinating link between sex, cellular plasticity, and the metabolic signaling pathways identified in our other components (Chittka et al., 2012; Yang et al., 2021). This theme of sex-differentiated plasticity was further exemplified by Component 278, which captured a female-biased shift toward a more precursor-like transcriptional state in the oligodendrocyte lineage and could have significant implications for myelin dynamics and repair capacity (Mathys et al., 2019; Akay et al., 2021; Lopez-Lee et al., 2024).

Our decomposition also resolved major axes of glial heterogeneity independent of sex. Component 177, for example, defined a purely microglial state characterized by the coordinated regulation of mitochondrial bioenergetics and autophagy/lysosomal pathways, adding further evidence to the mitochondria-microglia-inflammation axis Miao et al. (2025). This highlights a fundamental axis of variation in how microglia manage energy production and cellular quality control. In contrast, Component 87 revealed a different mode of glial crosstalk: a reciprocal relationship between astrocytes and OPCs linked to the expression of the immune checkpoint *VTCN1* and epigenetic poising of key glial regulatory kinases *AXL*, and *FLT4*, as well as calcium channel gene *CACNA1C*. This finding is particularly striking, as *CACNA1C* is a well-established risk gene for neuropsychiatric conditions, including mood disorders. Our observation of an epigenetic signature in the aging cortex aligns with recent findings, where chronic stress altered *Cacna1c* expression specifically in the prefrontal cortex of male mice (Bastos et al., 2024). This provides a powerful, cell-type-specific link to the long-standing calciumopathy hypothesis of AD (Stutzmann, 2007) and suggests that *CACNA1C* may be a critical node where vulnerability to both chronic stress and age-related pathology converge. Together, these components reveal distinct, uncoupled programs governing immunometabolism, cellular maintenance, and glial crosstalk in the aged brain.

These findings have significant therapeutic implications. The immunometabolic axis nominates mTOR and its upstream regulators as tractable entry points for modulating chronic neuroinflammation. The highly specific nature of the sex-dimorphic components, as well as the overall inter-individual variability in our cohort argue that a one-size-fits-all approach to treating neurodegenerative disease is likely to be insufficient. Precision therapeutics may need to be tailored to individuals e.g. boosting oligodendrocyte plasticity in females, or selectively modulating neuronal-stress pathways in sub-groups with heightened vulnerability. The replication of our key findings in the independent SEA-AD cohort (Gabitto et al., 2024) demonstrates that these programs are fundamental features of human brain aging, not artifacts of a single dataset. By unifying multiple molecular layers, our work moves beyond catalogues of single-omic changes to reveal the coherent biological themes that orchestrate cellular function in the aged brain, providing a framework for guiding the next generation of precision interventions.

### Limitations of the study

Our study has several limitations that provide context for our findings. First, there is an inherent imbalance in the depth and sample size of the integrated data modalities. The single-nucleus RNA-seq data, with its cellular resolution and largest sample cohort, necessarily provides the richest source of variation. In contrast, the proteomics data had the fewest features and samples. Consequently, while our approach integrates all layers, the resulting components are naturally weighted toward the more deeply characterized transcriptomic layer.

Second, this weighting extends to our validation strategy. The independent SEA-AD cohort provided an excellent resource for replicating the transcriptional signatures of our key components, strongly supporting their generalizability. However, as this dataset lacks matched multi-omic data, it provides indirect, rather than direct, validation for the specific epigenetic and proteomic patterns we identified.

Third, our results focus predominantly on glial-driven programs. This focus arose because the most robust multi-omic signals, those with coherent patterns across multiple molecular layers, were found in glial cell types. Our analysis did resolve neuron-specific components, many of which replicated known associations between neuronal gene expression and cognitive phenotypes. For example, we identified a purely inhibitory neuron component (Component 172) enriched for MAPK signaling and is associated with Alzheimer’s diagnosis (two-sided Mann-Whitney; *U* = 6726, *p* = 0.0004, FDR-adjusted *p* = 0.017) (Figure S6A-D) (Munoz & Ammit, 2010; Kim & Choi, 2010), and an excitatory neuron component (Component 277) enriched for GABAergic synapse genes and is associated with cognitive decline (Kruskal-Wallis; *H* = 21.9, *p* = 1.77 × 10^−5^, FDR-adjusted *p* = 0.002) (Figure S6E-H) (McQuail et al., 2015; Rozycka & Liguz-Lecznar, 2017). However, as these largely recapitulated findings from single-modality studies, we prioritized the novel, integrative glial programs that represent the unique strength of this multi-omic approach.

Finally, like all studies based on post-mortem tissue, our analysis provides a static snapshot of the aged brain. While we identify robust molecular states and their associations, we cannot directly capture the dynamic cellular transitions that occur over the course of aging and disease progression. Future work using longitudinal data or advanced model systems will be needed to dissect the causal relationships and temporal dynamics of the multi-omic programs identified here.

## METHODS

### Data sources

Data from the Religious Orders Study and Memory and Aging Project (ROSMAP) served as the primary dataset for this study (Bennett et al., 2018). ROSMAP consists of two longitudinal cohorts based at the Rush Alzheimer’s Disease Center (RADC). All participants enroll without known dementia and agree to annual evaluation and organ donation. Both cohorts were approved by the Institutional Review Board of Rush University Medical Center, and all participants signed informed-consent, data-repository, and Anatomical Gift Act documents.

Processed data modalities were downloaded from the AD Knowledge Portal^a^: single-nucleus RNA-sequencing (snRNA-seq) data (syn52293417) (Mathys et al., 2023); DNA methylation data (syn4896408) (De Jager et al., 2014); H3K9ac chromatin immunoprecipitation sequencing (ChIP-seq) data (syn3157275) (Klein et al., 2019); and mass-spectrometry proteomics data (syn17008935) (Ping et al., 2018). Our analysis focuses on 276 participants, well balanced for sex (147 female, 129 male), and spans the full range of cognitive status at death: 97 with no cognitive impairment, 65 with mild cognitive impairment, 3 with MCI plus another contributing condition, 92 with probable Alzheimer’s dementia, 12 with possible AD plus another condition, and 7 with other primary dementias. All molecular measurements (snRNA-seq, DNA methylation, H3K9ac ChIP-seq, and proteomics) were generated once per participant from a single dorsolateral prefrontal-cortex specimen, i.e. no technical replicates were included; the CERAD neuritic-plaque score was obtained from the same post-mortem tissue block, whereas cognitive status reflects the final antemortem clinical evaluation.

For independent validation of our transcriptomic findings, we used the “processed single cell data” from the Seattle Alzheimer’s Disease Brain Cell Atlas (SEA-AD) (Gabitto et al., 2024). Processed single-nucleus RNA-sequencing count matrices and associated metadata were downloaded from AWS as linked to from the Allen Brain Map portal^b^ for the dorsolateral prefrontal cortex.

For all datasets, we utilized the publicly available, processed data matrices to the greatest extent possible. Feature names (e.g., gene IDs, CpG loci identifiers) were retained from the original sources to ensure consistency. Minor pre-processing and harmonization steps required to integrate the different data types are detailed in the following sections.

### Data processing and harmonization for joint decomposition

All four data modalities were processed and harmonized to a final cohort of 276 individuals defined by the snRNA-seq dataset. Each matrix was aligned to have the same 276 sample columns in the same order, with missing data for any individual in any modality represented as `NaN` values, which the SDA model handles natively.

#### Single-Nucleus RNA-Sequencing (snRNA-seq)

Processing began with the pre-calculated AnnData object from the original ROSMAP study. After excluding two low-quality samples and all computationally predicted doublets, we filtered for samples containing at least 100 cells in each of the six major glial and neuronal cell types. For the final 276 samples, cell-type-specific pseudobulk profiles were generated by summing raw counts for each cell type. A gene filter was applied to retain genes with at least 10 counts in 10 or more pseudobulk samples. The final normalized expression data was structured as a three-dimensional tensor of (276 samples) × (6 cell types) × (25,098 genes), with no missing samples. The validation SEA-AD dataset was processed in a parallel manner, resulting in a tensor of (82 samples) × (10 cell types) × (31,648 genes).

#### H3K9ac ChIP-Sequencing (ChIP-seq)

The raw read count matrix was harmonized to match the projids of the 276-sample snRNA-seq cohort. Data was normalized to log2-transformed Counts Per Million (log2(CPM+1)). The final input matrix had dimensions of (276 samples) × (26,384 peaks), which included 83 samples with no ChIP-seq data (represented as missing values).

#### DNA methylation

We began with the ROSMAP imputed *β*-value matrix and harmonized it to the 276 individuals profiled by snRNA-seq. Each *β* value was temporarily binarized as methylated (1; *β* ≥ 0.5) or unmethylated (0; *β <* 0.5), preserving `NaN` for missing measurements. CpG sites that were invariant after binarization (all 0, all 1, or all missing) across the cohort were discarded, thereby removing constitutively methylated or unmethylated loci. The surviving probes retained their original continuous *β* values for downstream analysis, yielding a matrix of 51,798 variable CpG sites across 276 samples; 60 participants lacked methylation data..

#### Proteomics

We used the fully processed log2-transformed TMT proteomics data. After harmonizing sample identifiers to the 276 projids, we applied a stringent feature filter, removing any protein with a missing value in any of the present samples. The final input matrix had dimensions of (276 samples) × (5,230 proteins), with 171 participants lacked proteomic measurements.

### Tensor and matrix decomposition

To integrate the four processed omic datasets, we applied Sparse Decomposition of Arrays (SDA), a Bayesian framework for tensor and matrix decomposition (Hore et al., 2016). The snRNA-seq data was structured as a three-dimensional tensor of (276 samples) × (6 cell types) × (25,098 genes). The DNA methylation (51,798 CpG sites), H3K9ac ChIP-seq (26,384 peaks), and proteomics (5,230 proteins) data were each structured as two-dimensional matrices of (276 samples) × (features). These arrays were jointly decomposed by SDA, which is robust to the missing data present across the different modalities (Figure 1).

To ensure the identification of robust and reproducible biological signals, we performed ten independent runs of the SDA model. Each run was initiated with a request for 2,000 components and performed for 1,500 iterations. All non-zero components from all ten runs were then pooled and subjected to a rigorous clustering and filtering procedure. Following the established protocol (Hore et al., 2016), we performed hierarchical clustering on the components based on their pairwise dissimilarity, defined as one minus the absolute Pearson correlation of their sample loading scores. Flat clusters were formed using a cophenetic distance threshold of 0.4. A stringent stability criterion was then applied, retaining only those clusters containing components from at least five of the ten independent runs. This process yielded a final set of 546 stable components. A final, representative loading score for each feature and sample was calculated by averaging the scores of all components within each stable cluster, after aligning their signs.

A key feature of the SDA framework is its ability to perform automatic feature selection through its use of a spike-and-slab prior. For each feature’s loading score in a given component, the model calculates a Posterior Inclusion Probability (PIP), a value between 0 and 1 representing the confidence that the feature is genuinely associated with that component. For all downstream biological interpretation, we applied a stringent threshold, considering only features with a PIP > 0.5 to be robustly associated with a component.

This step is crucial for two reasons. First, it creates a sparse representation, effectively reducing the vast number of initial features to a core set of high-confidence drivers for each biological program. Second, it means that components are not required to be active across all modalities. A significant implication of this feature selection is that many of the resulting components are driven by only one or two omic types (e.g., a purely transcriptomic signature or a coupled transcriptomic-epigenomic axis), allowing the model to capture both single-modality and truly integrative multi-omic patterns. The final loading scores reported for these selected, high-confidence features represent the expected value from the model’s posterior distribution.

The interpretation of each component’s effect is based on the multiplicative relationship between its loading scores. For the three-dimensional snRNA-seq data, a component’s contribution to the expression of a given gene in a particular cell type for a specific sample is proportional to the product of the three corresponding loading scores (sample, cell type, and gene). Consequently, the sign of this product determines the direction of the effect: a positive product indicates that the component contributes to higher relative expression, while a negative product indicates lower relative expression. The magnitude of this contribution is, in turn, determined by the product of the magnitudes of these scores. This principle extends to the two-dimensional matrices, where a component’s effect is the product of the sample and feature loading scores.

#### Validation in an independent cohort

To assess reproducibility, we correlated each ROSMAP component with components derived from the SEA-AD snRNA-seq dataset (dorsolateral prefrontal cortex). For a given ROSMAP component we first extracted genes with PIP > 0.5, intersected this set with the SEA-AD gene list, and assembled two loading vectors (ROSMAP vs. SEA-AD) on the common genes. We then computed the Spearman correlation coefficient between every pair of vectors and recorded, for each ROSMAP component, the SEA-AD component that achieved the highest *absolute* correlation. Absolute correlation was used because the sign of SDA loadings is arbitrary: multiplying all sample and feature loadings by −1 yields an equivalent factor and leaves the reconstructed data unchanged.

#### Component prioritization

From the 546 ROSMAP components we selected those most likely to represent robust, integrative biology through a two-step filter.

1. Multi-omic coherence. We required high-confidence features (PIP > 0.5) in at least three of the four modalities (gene expression, DNA methylation, H3K9ac peaks, and proteomics) thereby focusing on genuinely integrative axes.
2. Cross-cohort reproducibility. Among these coherent components we retained only those whose transcriptional signature correlated strongly with a SEA-AD component (|r| > 0.4 by Spearman). Because of the sign indeterminacy noted above, the magnitude of the correlation, not its direction, indicates biological concordance.

The resulting set of integrative, reproducible components forms the basis of the detailed analyses presented in the Results.

#### Genetic association and eQTL analysis

To identify genetic variants driving component scores, we first performed a genome-wide association study (GWAS) for each of the 546 components, using the sample loading scores as a quantitative trait. This was performed using PLINK 2.0 with a linear regression model, including age, sex, and the first ten genetic principal components as covariates. SNPs passing a standard genome-wide significance threshold (*p <* 1 × 10^−15^) were considered for downstream analysis.

For each component with at least one significant SNP, we performed a cell-type-specific expression quantitative trait locus (eQTL) analysis to test the association between the SNP genotype and the expression of high-confidence genes (PIP > 0.5) from that component. This association was modeled using an ordinary least squares regression followed by an F-test (equivalent to ANOVA) (`expression ∼ C(genotype)`) and was conducted separately for each of the six major cell types using their respective pseudobulk expression profiles. The resulting eQTL effects were visualized using the boxplots and stripplots shown in the figures.

#### Association with phenotypes

For each component we tested whether sample loading scores differed across key clinical variables. Binary traits, biological sex (129 males, 147 females) and neuropathological diagnosis (170 CERAD-AD vs. 106 non-AD), were evaluated with two-sided Mann-Whitney U tests.

To examine cognition, we focused on a three-level ordinal scale capturing the canonical clinical continuum while excluding mixed or non-AD dementias. Participants were grouped as: (i) no cognitive impairment (n = 97); (ii) mild cognitive impairment without other contributing pathology (n = 65); and (iii) probable Alzheimer’s dementia with no additional primary aetiology (n = 92). Cases with comorbid contributions to impairment (n = 15) or other primary dementias (n = 7) were excluded from this analysis. No additional covariates (e.g., age at death, postmortem interval, RNA integrity) were included in these univariate tests. Differences across the three ordered groups were assessed with Kruskal-Wallis test.

P-values for all 546 components were corrected for multiple testing within each phenotype using the Benjamini-Hochberg false-discovery rate (FDR); components with FDR < 0.05 were considered significant.

#### Pathway enrichment analysis

To identify enriched biological pathways for each component, we used the gseapy Python package, an interface to the Enrichr toolset (Xie et al., 2021). For a given component, we selected all features (genes, proteins, etc.) with a Posterior Inclusion Probability (PIP) > 0.5. The official gene symbols corresponding to these high-confidence features were used as the input gene list for enrichment analysis against the KEGG 2021 Human gene set. Pathways with an FDR-adjusted p-value < 0.05 were considered significantly enriched. The most significant pathway per component per omic type is reported in Supplementary Table 1.

#### Motif enrichment analysis

For each component, we searched for transcription-factorbinding motifs enriched in its H3K9ac peaks using the HOMER suite (v5.1). Genomic coordinates of all peaks with PIP >0.5 were exported as a BED file and passed to `findMotifsGenome.pl` (default parameters) against the GRCh37.p13 reference genome. HOMER builds a GC-matched background set automatically. Motifs with a HOMER-reported enrichment *p* < 0.05 (binomial test) were deemed significant; the most significant motif per component is reported in Supplementary Table 1.

#### Plotting and visualization

All figures in this manuscript were generated in Python. Data manipulation and analysis were performed using the *pandas* and *scipy* libraries. Plots were generated using *matplotlib* and *seaborn*. The *statannotations* package was used to display statistical significance on box plots, and the *adjustText* package was used for non-overlapping label placement in scatter plots. To effectively visualize sample loading plots containing extreme outliers, we defined outliers as any data point more than 30 times the interquartile range (IQR) from the upper or lower quartile. For these specific plots, the *brokenaxes* package was used to create a discontinuous y-axis, allowing for the clear display of both the main data distribution and the extreme outlier values on the same figure, or not included in the boxplots.

## ACKNOWLEDGMENTS

We thank Dr. Robert Marr for his insightful feedback during the preparation of this manuscript. The results published here are in whole or in part based on data obtained from the Rush Alzheimer’s Disease Center, Rush University Medical Center, Chicago, and University of Washington Alzheimer’s Disease Research Center. We are deeply grateful to the participants of the Religious Orders Study and Memory and Aging Project (ROSMAP), as well as the Seattle Alzheimer’s Disease Brain Cell Atlas (SEA-AD) consortium for their invaluable contributions to this research.

This work was supported by grants from the National Institutes of Health (NIH), National Institute on Aging (NIA) and the Alzheimer’s Association (AA). Specific funding for the authors includes: NIH grants NS135301 and NS135306 to D.A.P.; NIA grant 5R00AG059953-05 and AA grant 24AARG-1188074 to H.C.H.; and NIH grants R01AG065628 and R21AG083638 to G.E.S. ROSMAP is supported by NIH grants P30AG10161, P30AG72975, R01AG17917, R01AG015819, U01AG072572, and U01AG046152.

## AUTHOR CONTRIBUTIONS

J.P.W. conceptualized the study, processed the data, performed the formal analysis and visualization, and wrote the original draft of the manuscript. H.C.H., D.A.B., M.L., D.A.P., and G.E.S. contributed to study design, provided critical feedback, and aided in the interpretation of results. All authors reviewed and edited the manuscript and approved the final submission.

## DATA AVAILABILITY

All primary data analysed in this study are publicly available. The discovery multi-omic datasets (snRNA-seq, DNA methylation, H3K9ac ChIP-seq and TMT proteomics) were obtained from the Religious Orders Study and Memory and Aging Project (ROSMAP) through the AD Knowledge Portal under accessions syn52293417, syn4896408, syn3157275 and syn17008935, respectively. Access to ROSMAP files requires a free Synapse account and acceptance of the ROSMAP data-use agreement; no other restrictions apply. Validation was performed with the Seattle Alzheimer’s Disease Brain Cell Atlas (SEA-AD) single-nucleus RNA-seq dataset, available without restriction via the Allen Brain Map portal and Amazon S3. All derived data products generated here; including the complete set of 546 component loading matrices, figure source data and Supplementary Table 1; have been deposited on either Synapse with the ROSMAP primary data or Zenodo by publication. No additional datasets were used in this study.

## List of Supplementary Tables

S1 **Legend for Supplementary Table 1: Comprehensive annotation of 546 multi-omic components**.

## List of Supplementary Figures

S1 **Component 21 showing ectopic olfactory receptor expression and its genetic regulation**.

S2 **Cross-validation of the microglial gene expression (Component 351)**. *Related to Figure 2 in the Main Text*

S3 **Replication of the microglial-astrocytic axis captured by Component 2**. *Related to Figure 3 in the Main Text*

S4 **A sex-biased component, 278, captures a balance between OPC and mature oligodendrocyte states**. *Related to Figure 4 in the Main Text*

S5 **A microglial bioenergetic and quality control axis (Component 177)**. *Related to Figure 5 in the Main Text*

S6 **Neuron-specific components, 172 and 277, replicate known associations with Alzheimer’s disease pathology and cognitive decline**.

**Table S1. Legend for Supplementary Table 1: Comprehensive annotation of 546 multi-omic components**.

**Supplementary Table 1 is provided as a separate file (**Table_S1.csv**). The descriptions below detail the content of each column in that file**.

`Component` The unique identifier for each of the 546 components.

`omics_count` The number of omic modalities (out of four) with at least one high-confidence feature (Posterior Inclusion Probability [PIP] > 0.5) for that component.

`Top_positive_cell_type, Top_negative_cell_type` The cell types with the highest positive and negative loading scores, respectively, from the snRNA-seq data. A blank cell indicates that no RNA features passed the PIP > 0.5 threshold for that component.

### Clinical Phenotype Associations

`Sex, ceradsc, cogdx` For each phenotype, the table reports the Benjamini-Hochberg adjusted p-value (FDR). A blank cell indicates the association was not significant (FDR ≥ 0.05). The specific statistical tests and variable coding were as follows:

- `Sex:` p-value from a two-sided Mann-Whitney U test comparing Male vs. Female.
- `ceradsc:` p-value from a two-sided Mann-Whitney U test comparing groups based on the CERAD score for neuritic plaques. Scores 1-2 were grouped as “Alzheimer’s Disease” and scores 3-4 were grouped as “No”.
- `cogdx:` p-value from a one-way Kruskal-Wallis test comparing three groups based on the final cognitive diagnosis: “No cognitive impairment” (score 1), “Mild cognitive impairment” (score 2), and “Alzheimer’s Disease” (score 4).

`SEA-AD_comp, SEA-AD_corr` The component index from the SEA-AD dataset that showed the highest absolute correlation with the ROSMAP component’s transcriptional signature, and the value of that Spearman correlation coefficient.

### Top Features and Pathways by Omic Type

`Top_Positive/[omic], Top_Negative/[omic]` The single feature (e.g., gene, CpG site, protein, or peak) with the highest positive or lowest negative loading score, respectively, among all features with a PIP > 0.5 for that modality. A blank cell indicates that no features for that omic type passed the PIP > 0.5 threshold for that component.

`GeneExpression_Pathway, CpG_Site_Pathway, Protein_Pathway` The top enriched KEGG pathway identified by Enrichr from the set of high-confidence features (PIP > 0.5) for that modality. A blank cell indicates that no features for that omic type passed the PIP > 0.5 threshold, and no enrichment analysis was performed.

`ChIP_Motif` The top enriched transcription factor binding motif identified by HOMER from the set of high-confidence ChIP-seq peaks (PIP > 0.5). A blank cell indicates that no ChIP-seq peaks passed the PIP > 0.5 threshold, and no motif analysis was performed.

**Figure S1.**
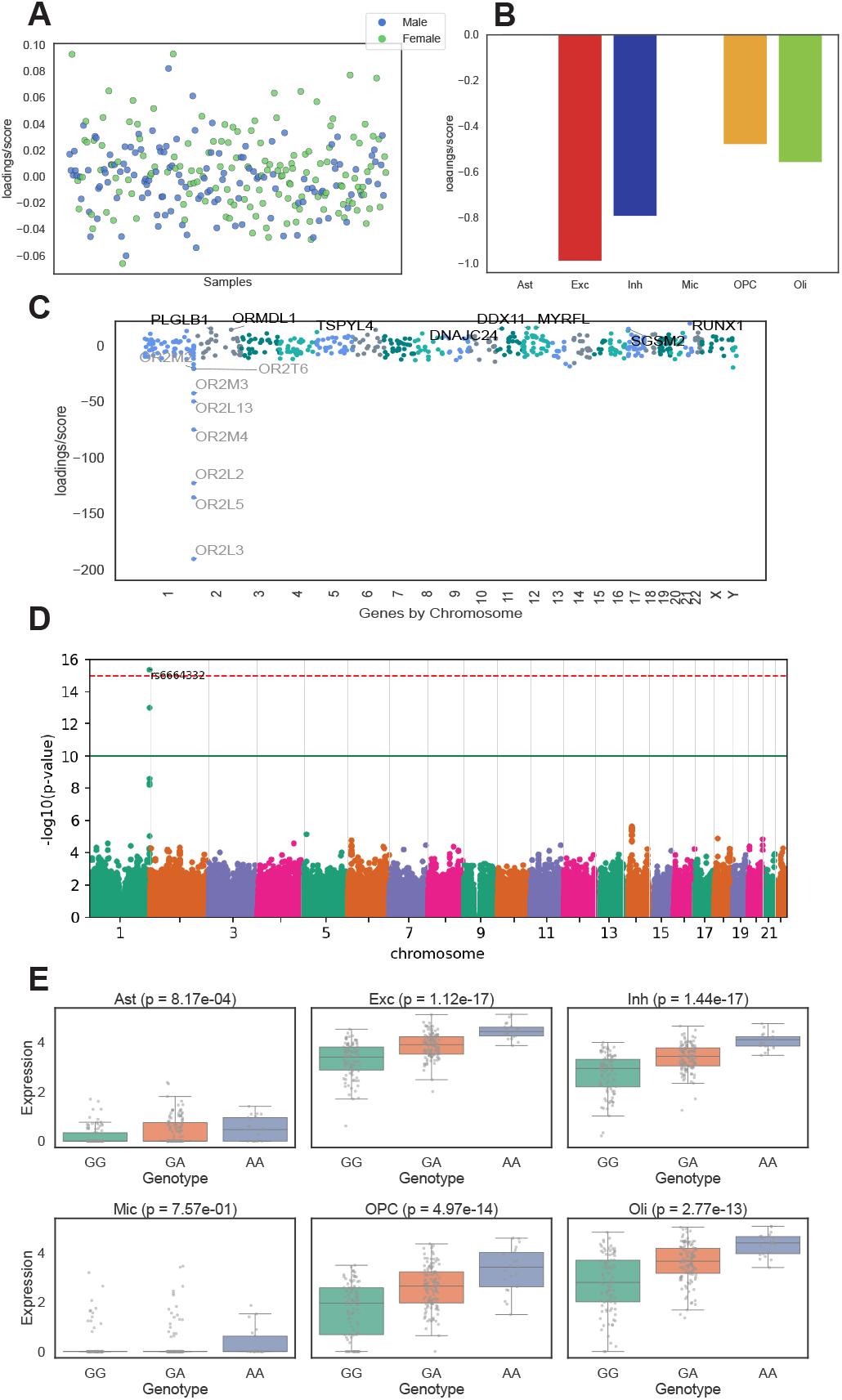
Component 21 showing ectopic olfactory receptor expression and its genetic regulation. **(A)** Sample-level loading scores for Component 21 across individuals in the ROSMAP dataset, colored by reported sex. **(B)** Cell-type loading scores, indicating that Component 21 is primarily driven by expression in inhibitory (Inh) and excitatory (Exc) neurons, oligodendrocytes (Oli), and oligodendrocyte precursor cells (OPC), with minimal contributions from astrocytes (Ast) and microglia (Mic). **(C)** Gene loading scores plotted by chromosomal position, highlighting a striking enrichment for olfactory receptor (OR) genes on chromosome 1. Top-weighted genes include *OR2L3, OR2L5, OR2L2, OR2L13, OR2M4*, and others within the OR gene cluster. **(D)** Genome-wide association analysis, using Component 21 sample loading scores as a quantitative phenotype, identifies a significant locus on chromosome 1 centered on SNP rs6664332 (green box). **(E)** Expression quantitative trait locus (eQTL) analysis for *OR2L3* stratified by rs6664332 genotype, illustrated by box plots showing median ± IQR; whiskers extend to 1.5 × IQR. A strong genotype-dependent expression pattern is observed in both neuronal cells; Exc and Inh, with similar effects seen in the glia; Oli and OPC, but not Ast and Mic (p-values from an ordinary least squares ANOVA).

**Figure S2.**
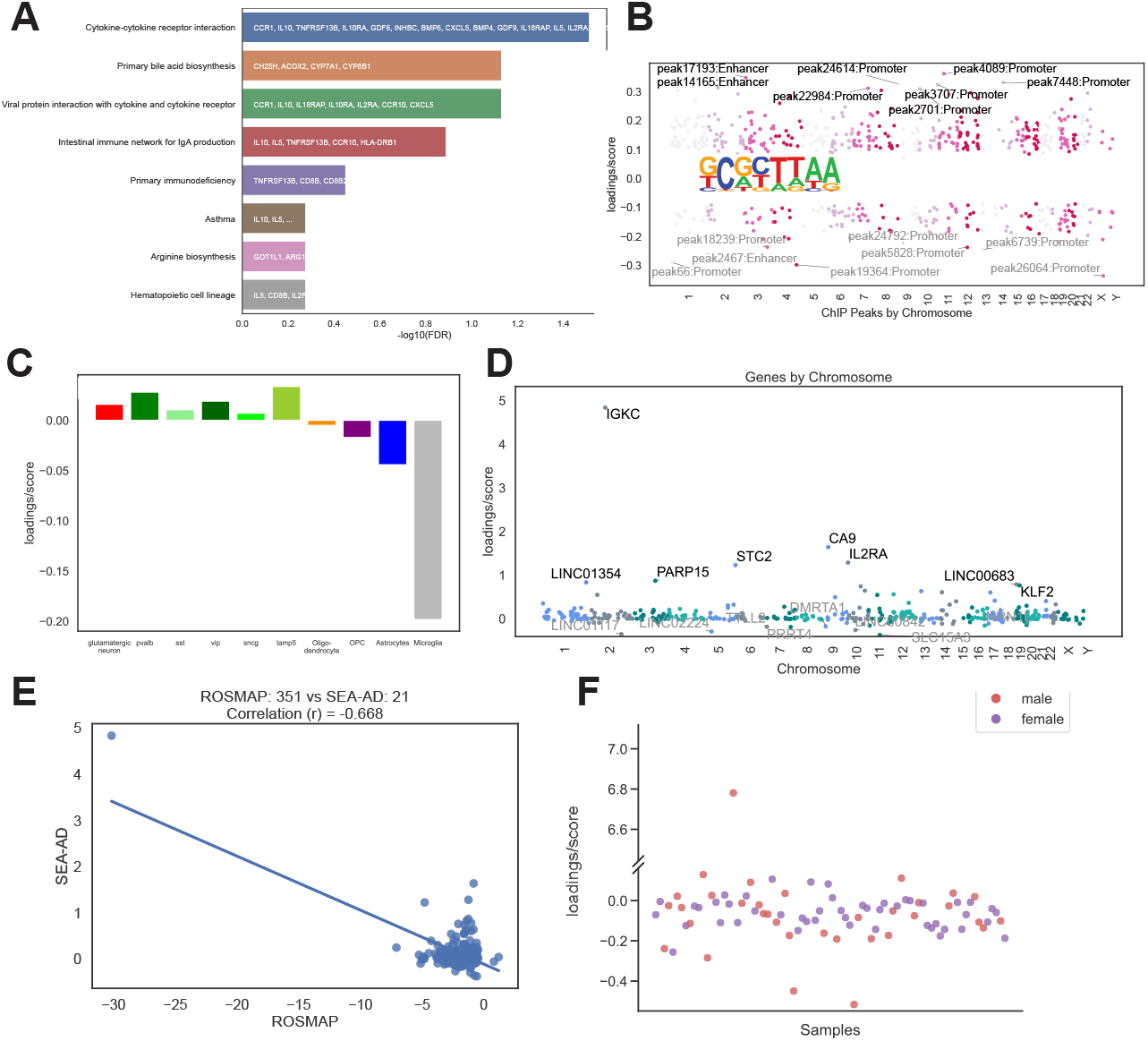
Cross-validation of the microglial gene expression (Component 351). *Related to Figure 2 in the Main Text* **(A)** KEGG pathways enriched for Component 351 genes, primarily immune and cytokine related. **(B)** H3K9ac ChIP-seq loadings highlight promoter and enhancer peaks; positively loaded peaks share a *BARHL1*-like homeobox motif, suggesting developmental regulators are re-engaged under inflammatory stress. **(C)** In the SEA-AD cohort, Component 21 shows a matching cell-type pattern with dominant microglial loadings. **(D)** The same SEA-AD component reproduces the ROSMAP gene signature, including *IGKC, HLA-DRB1*, and other immune transcripts. **(E)** Gene-loading scores in SEA-AD versus ROSMAP show a high Spearman’s correlation coefficient, confirming cross-cohort robustness. **(F)** SEA-AD sample scores mirror the ROSMAP distribution, with a single outlier exhibiting markedly elevated loading, consistent with focal inflammatory activation.

**Figure S3.**
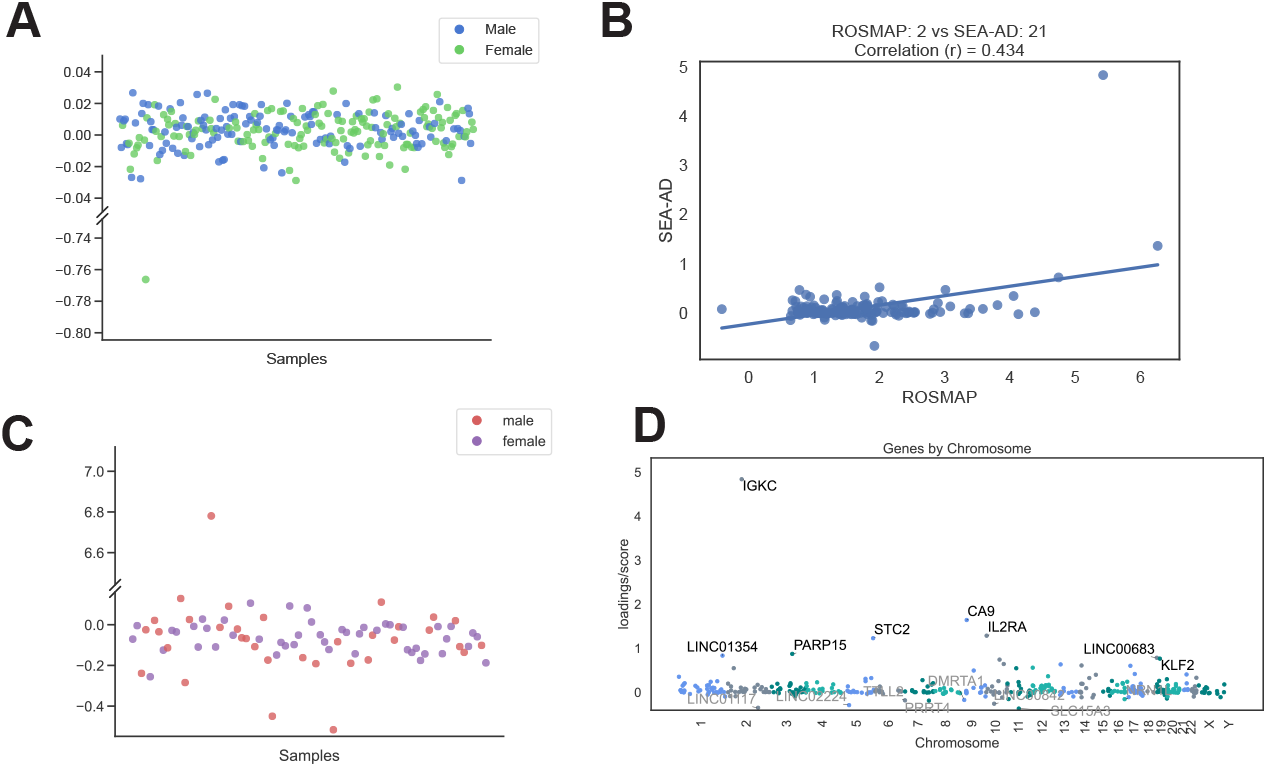
Replication of the microglial-astrocytic axis captured by Component 2. *Related to Figure 3 in the Main Text* **(A)** ROSMAP sample scores for Component 2 show broad inter-individual spread with a small set of high-scoring outliers. **(B)** Gene-loading correlation between ROSMAP Component 2 and the matched SEA-AD component 21 (with Spearman’s correlation coefficient) confirms cross-cohort reproducibility of the transcript signature. **(C)** SEA-AD component 21 sample scores exhibit a similarly skewed distribution, including an extreme outlier mirroring the ROSMAP distribution. **(D)** Gene loadings in SEA-AD component 21 recapitulate the immunoglobulin-rich microglial signature led by *IGKC*, along with several uncharacterized regulatory transcripts, reinforcing the shared immune-glial program.

**Figure S4.**
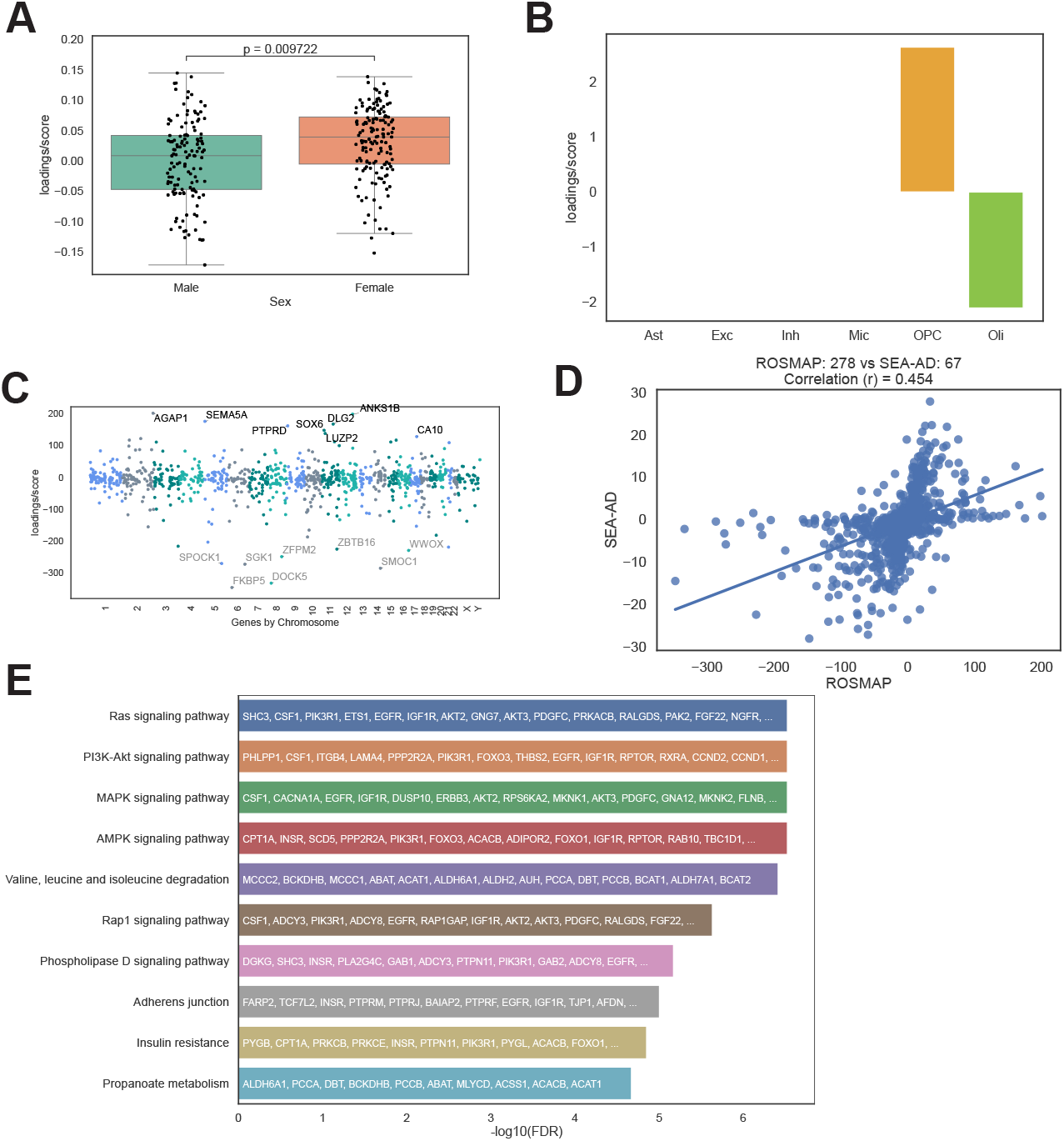
A sex-biased component, 278, captures a balance between OPC and mature oligodendrocyte states. *Related to Figure 4 in the Main Text* **(A)** Box plot (median ± IQR; whiskers extend to 1.5 × IQR) showing sample scores for Component 278, separated by sex. Female scores are significantly higher (FDR-adjusted *p* = 0.0097). **(B)** Cell-type loading scores for Component 278, showing a strong positive contribution from oligodendrocyte precursor cells (OPCs) and a strong negative contribution from mature oligodendrocytes (Oli). **(C)** Gene expression loadings for Component 278. Top-weighted genes include the oligodendrocyte differentiation factor *SOX6* and the stress-response gene *FKBP5*. **(D)** Scatter plot showing a significant positive correlation between ROSMAP Component 278 and a corresponding component in the SEA-AD dataset, demonstrating cross-cohort replicability with Spearman’s correlation coefficient shown. **(E)** KEGG pathway enrichment analysis for genes driving Component 278. Top enriched KEGG pathways include Ras, PI3K-Akt, and MAPK signaling, which are critical for cell growth and differentiation.

**Figure S5.**
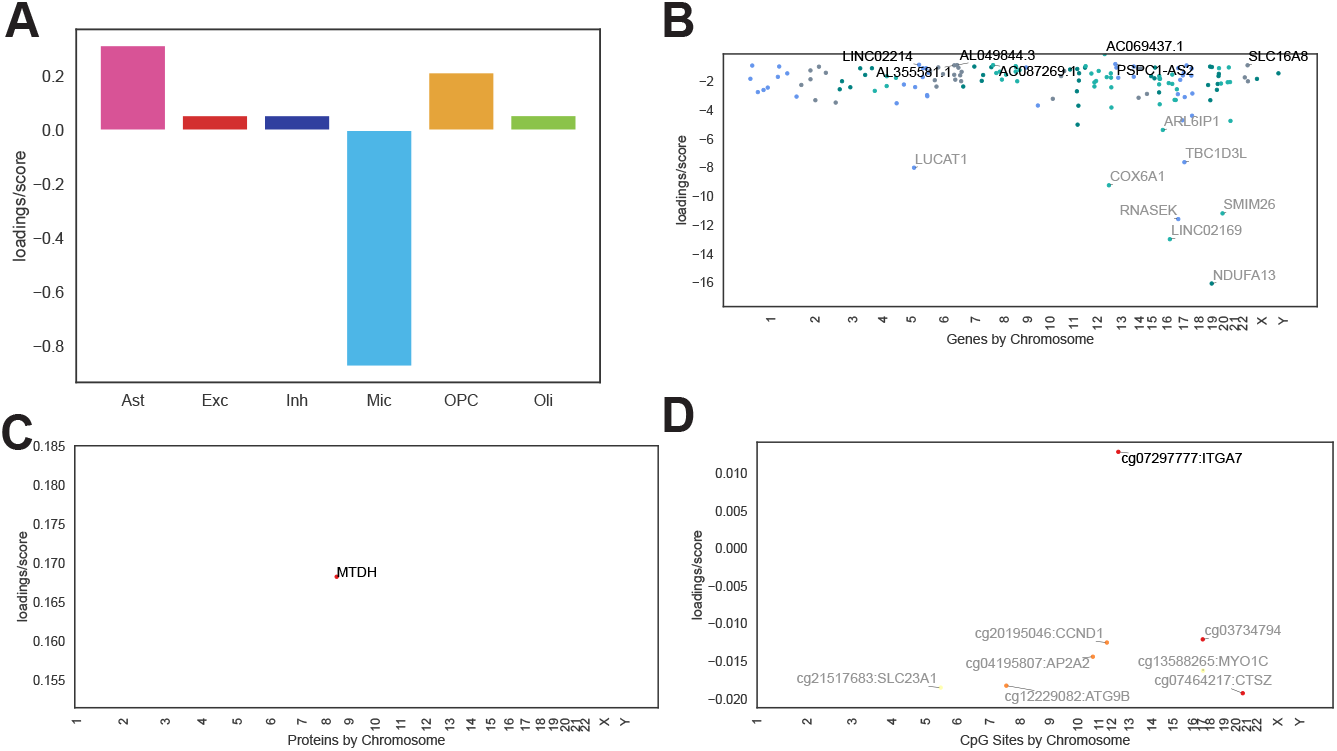
A microglial bioenergetic and quality control axis (Component 177). *Related to Figure 5 in the Main Text* **(A)** Cell-type loadings show the component is driven almost exclusively by microglia (Mic). **(B)** Gene expression loadings are defined by genes involved in mitochondrial bioenergetics, including the Complex I subunit *NDUFA13* and the Complex IV subunit *COX6A1*. **(C)** The top protein loading for the component is Metadherin (MTDH). **(D)** DNA methylation loadings highlight a coordinated signature including key autophagy genes (*ATG9B, CTSZ*) and the oppositely-regulated cell adhesion gene *ITGA7*.

**Figure S6.**
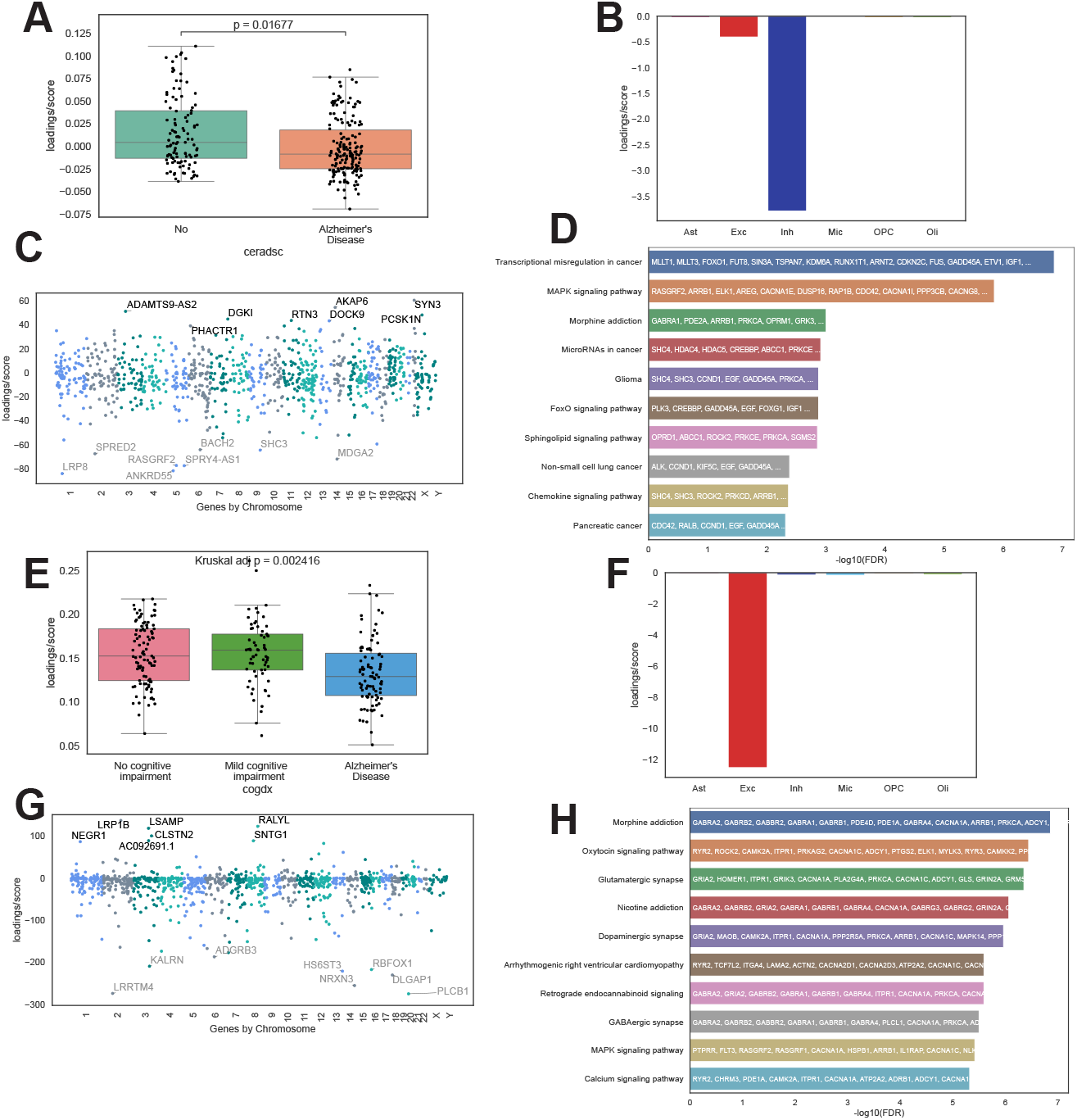
Neuron-specific components, 172 and 277, replicate known associations with Alzheimer’s disease pathology and cognitive decline. **(A)** Box plot (median ± IQR; whiskers extend to 1.5 × IQR) of sample scores for Component 172, showing a significant association with CERAD score for Alzheimer’s disease (FDR-adjusted *p* = 0.017). **(B)** Cell-type loadings for Component 172, indicating the signal is driven almost exclusively by inhibitory neurons. **(C)** Gene expression loadings for Component 172. **(D)** KEGG pathway enrichment for Component 172, showing enrichment for MAPK signaling. **(E)** Box plot (median ± IQR; whiskers extend to 1.5 × IQR) of sample scores for Component 277, showing a significant association with cognitive decline (FDR-adjusted *p* = 0.002). **(F)** Cell-type loadings for Component 277, indicating the signal is driven overwhelmingly by excitatory neurons. **(G)** Gene expression loadings for Component 277. **(H)** KEGG pathway enrichment for Component 277, showing strong enrichment for synaptic pathways, including GABAergic and glutamatergic synapses.

## Footnotes

a https://adknowledgeportal.synapse.org

b https://registry.opendata.aws/allen-sea-ad-atlas/

